# Small RNA-Mediated Metabolic Reprogramming Enables Ciprofloxacin Utilization in *Klebsiella* sp. SG01

**DOI:** 10.1101/2025.07.11.664498

**Authors:** Sriradha Ganguli, Ranadhir Chakraborty

## Abstract

Small regulatory RNAs (sRNAs) play a key role in the complex regulatory networks that bacteria use to adapt to antibiotic stress. This study clarifies how sRNAs help *Klebsiella* sp. SG01 metabolically adapts to ciprofloxacin (CIP). Nine potential sRNAs that target important metabolic and stress-response pathways were identified by transcriptomic profiling, including *glmZ*, *sgrS*, *gcvB*, and *spot42*, as well as an FMN riboswitch. TargetRNA3 and IntaRNA functional annotation predicted regulatory effects on TCA cycle flux, redox balance, and the transport of sugars and amino acids. Interestingly, repression of CRP and upregulation of *spot42* imply that byproducts of CIP degradation resemble a glucose-rich state. This conjecture was supported by metabolomics, which verified the buildup of short-chain acids and purine intermediates. A change in regulation was also noted, with ProQ and CsrA levels rising in tandem with Hfq’s downregulation. Our knowledge of microbial metabolic plasticity under xenobiotic pressure has been expanded by these findings, which collectively reveal a novel post-transcriptional mechanism that permits antibiotic assimilation and stress resilience in bacteria.

## 1. Introduction

The classical model of gene expression by Crick (1958, 1970) speaks of a linear flow of genetic information from DNA transcribed to RNA which is translated to proteins. The discovery of noncoding RNAs (ncRNAs), specifically small regulatory RNAs (sRNAs) ranging between 50-500 nucleotides (nt) long transformed the idea of regulation of gene expression across all domains of life (Cech and Steitz, 2014; Pennisi, 2010).

Bacteria are subjected to massive environmental perturbations and their survival or adaptation is controlled at every level-transcriptional, translational and post-transcriptional. Hence, along with transcription factors, sRNAs are potent players of post-transcriptional modulation in bacteria (Mizuno et al. 1984; Sorek and Cossart, 2010). They mostly act by base-pairing with target mRNAs by either stabilizing the mRNAs or inhibiting translation initiation. RNA chaperones like Hfq, ProQ and CsrA participate in such interactions to enhance the process (Holmqvist et al. 2018; Hor et al. 2020).

Heterotrophic bacteria, including *Klebsiella*, have robust metabolic flexibility which allows them to survive using a wide variety of carbon sources. Previously, transcriptional factors such as LacI were thought to be responsible for efficient substrate utilisation (Jacob and Monod, 1961) but modulation by sRNAs play a vital role when it comes to metabolic reprogramming under oxidative, nutritional or cell envelope stress. Little has been known about sRNA regulation under xenobiotic stress such as antibiotics. The recalcitrant antibiotic ciprofloxacin (CIP) which is frequently detected in the environment worldwide causes oxidative and genotoxic stress to the residing microbes, and hence does sets off intricate regulatory cascades in them

The metabolic adaptation of *Klebsiella* sp. SG01 to ciprofloxacin as its only carbon source is examined in this work through sRNA. Using transcriptomics, computational tools, and metabolomics, we find new and known sRNAs that are enriched under antibiotic stress and investigate their predicted targets and regulatory pathways. By analyzing this regulatory environment, we anticipate to reveal how bacteria use antibiotics as metabolic substrates as well as stressors, illuminating a new paradigm of microbial adaptation and biotransformation.

## 2. Results

### 2.1. Identification of Candidate sRNAs

The intergenic regions of *Klebsiella* sp. SG01 were searched for putative small RNAs (sRNAs) candidates. There were a total of 529 sequences, with 107 sequences in the 15-50 nt range, 299 into the 50-250 nt range, and 123 into the 200-500 nt range. These extracted sequences were compared against known non-coding RNAs in the Rfam database (Ontiveros-Palacios et al. 2024). Thirty-one candidate non-coding RNAs were initially found from this screening. Ten regulatory RNA elements were finally selected after additional filtering according to transcriptomic expression levels, structural characteristics, and functional significance. These comprised one riboswitch, the FMN aptamer (RFN element), and nine sRNAs: *glmZ*, *sgrS*, *gcvB*, *spot42*, *micA*, *istR*, *sraL*, *icsR*, and *fnrS*. These ten sRNAs were further evaluated to study their role in metabolising CIP by the strain SG01.

### 2.2. Length and GC Content Distribution of Identified sRNAs

Prokaryotic sRNAs are typically 50–500 nucleotides (nt) long, so we excluded sequences shorter than 15 nt or longer than 500 nt. Eight candidates fell in this range (50-200nt mostly) while two of the regulatory RNA elements were longer than 200 nt.

Three sRNAs had less than 50% GC content, according to GC content analysis, whereas the remaining six sRNAs and the FMN aptamer had GC content of at least 50%. The FMN riboswitch was also structurally stable with h a GC content of 59% and a length of 153 bp, typically found in regulatory RNAs.

Sequence length and GC content are two vital characteristics that gives us information about the possible folding dynamics and stability of sRNAs under stress (Fig. 1).

**Figure 1.**
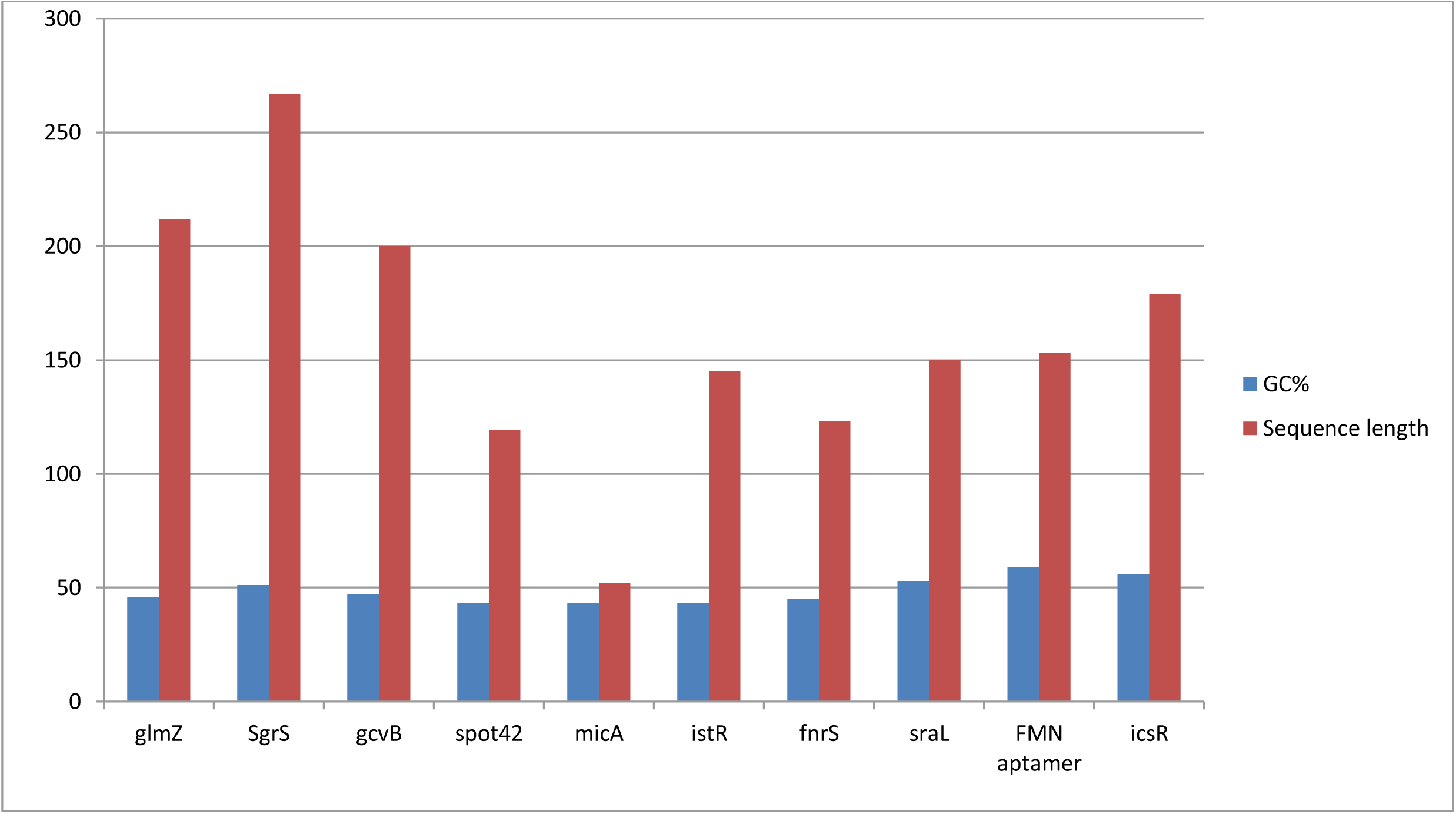
GC content and sequence length distribution of Identified sRNAs. The figure illustrates the distribution of GC content (%) relative to the sequence length (in nucleotides) for the identified sRNAs. The observed variability in GC composition across different sRNA lengths offers insights into their structural stability and potential regulatory functions.

### 2.3. Secondary structure and Functional analysis of sRNAs

RNAfold was used to predict the secondary structures of the identified sRNAs (Lorenz et al. 2011). The corresponding minimum free energy (MFE) values, which indicate thermodynamically stable configurations, ranged from -20 to -80 kcal/mol (Garcia-Martin et al. 2015). The structural characteristics, genomic settings, and anticipated stability of these sRNAs are compiled in Table 1.

**Table 1.**
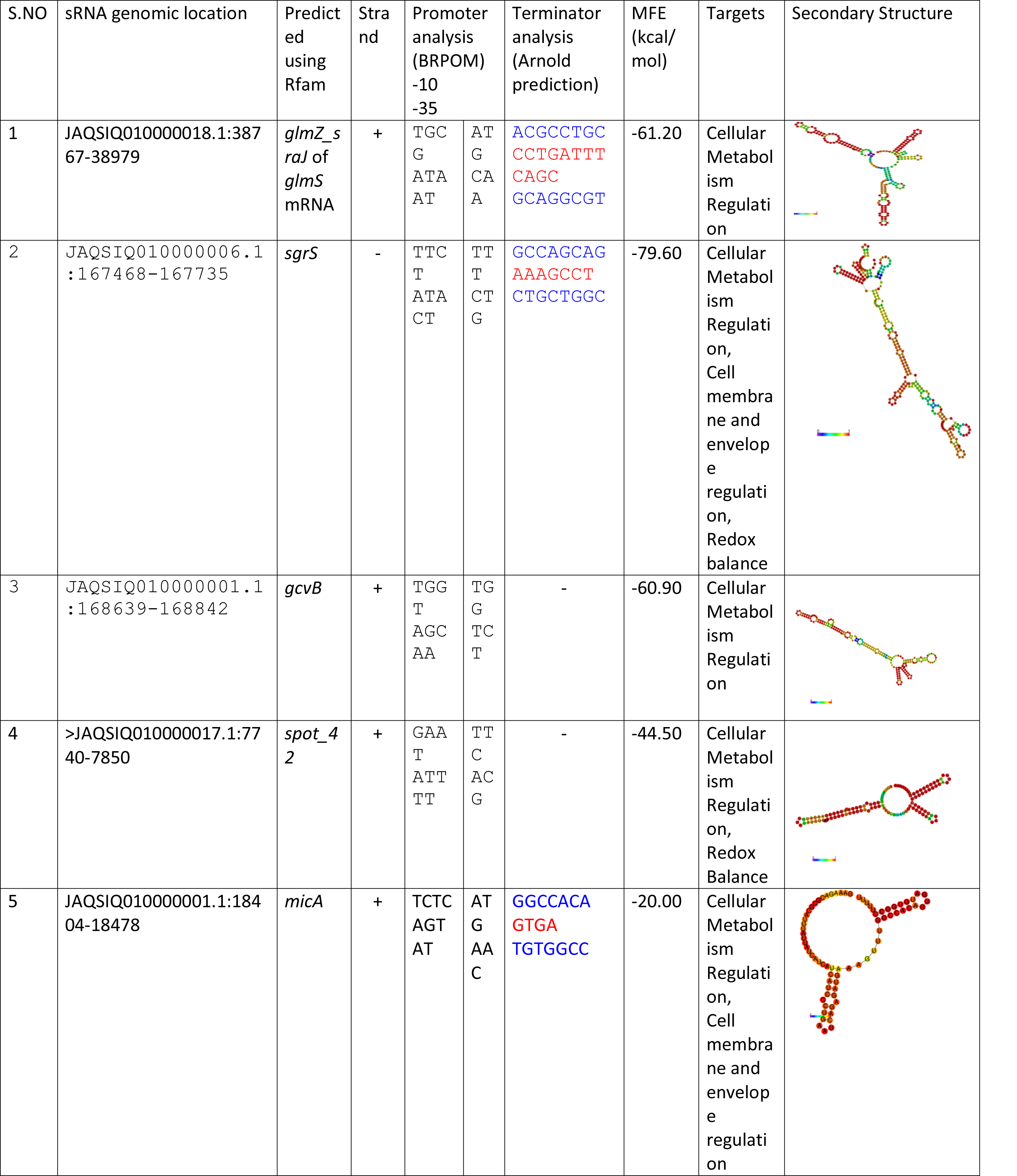
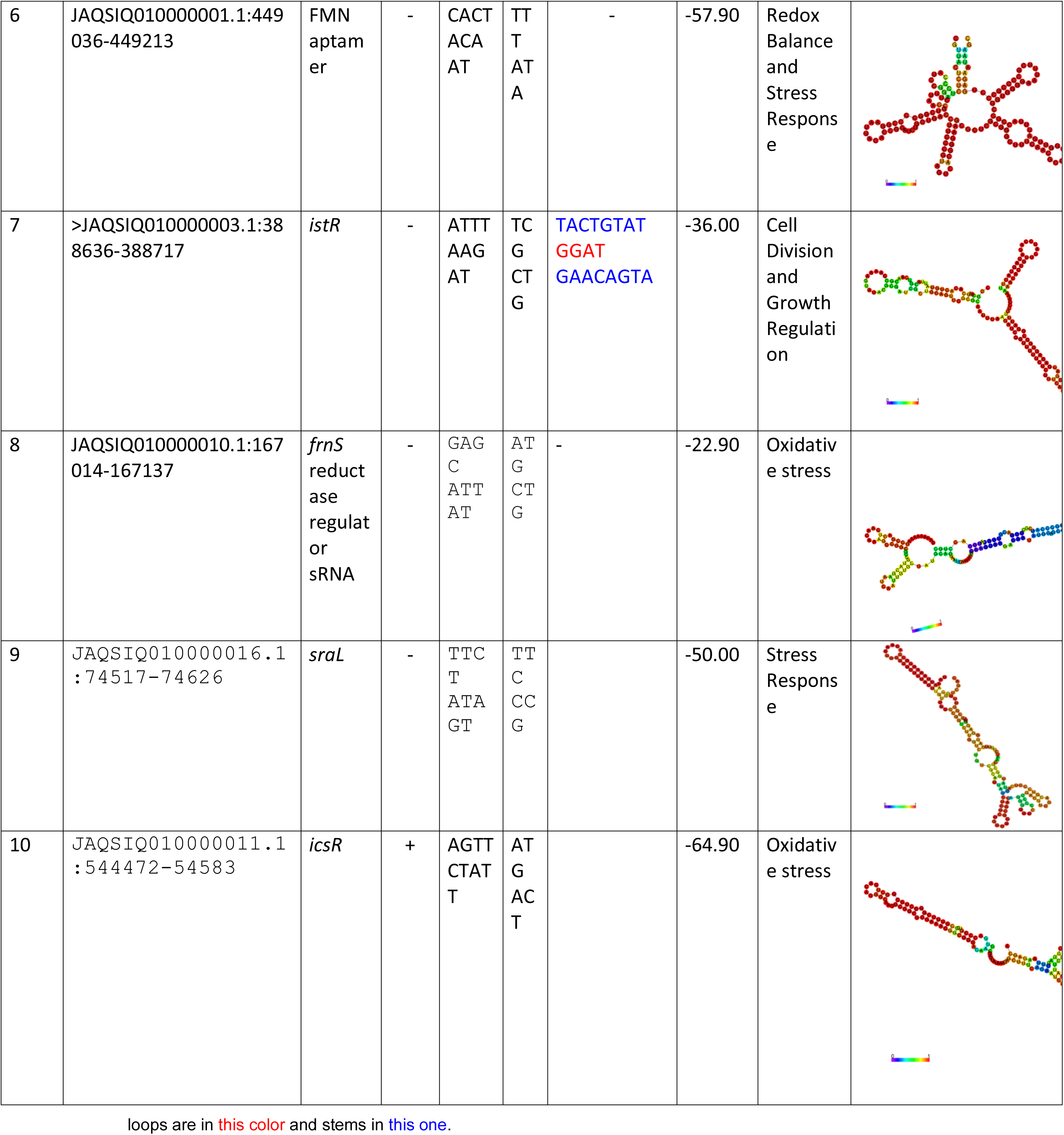
Genomic organization and structural features of small regulatory RNAs (sRNAs) identified from transcriptome data in *Klebsiella* sp. SG01 under ciprofloxacin (CIP) stress. This table provides detailed information on the genomic location, strand orientation (+/−), presence of promoter and terminator elements, and minimum free energy (MFE) values derived from predicted secondary structures of the identified sRNAs.

We predicted mRNA targets using TargetRNA3 in order to investigate the regulatory functions of these sRNAs during ciprofloxacin (CIP) assimilation. Experimental datasets from RIL-seq (Melamed et al. 2016), MAPS (Lalaouna et al. 2015), and CLASH (Waters et al. 2017) are integrated into this computational platform. A number of sRNAs were linked to central metabolic and stress-response pathways, including *glmZ*, *sgrS*, *gcvB*, *spot42*, and *micA* (Fig. 2).

**Figure 2.**
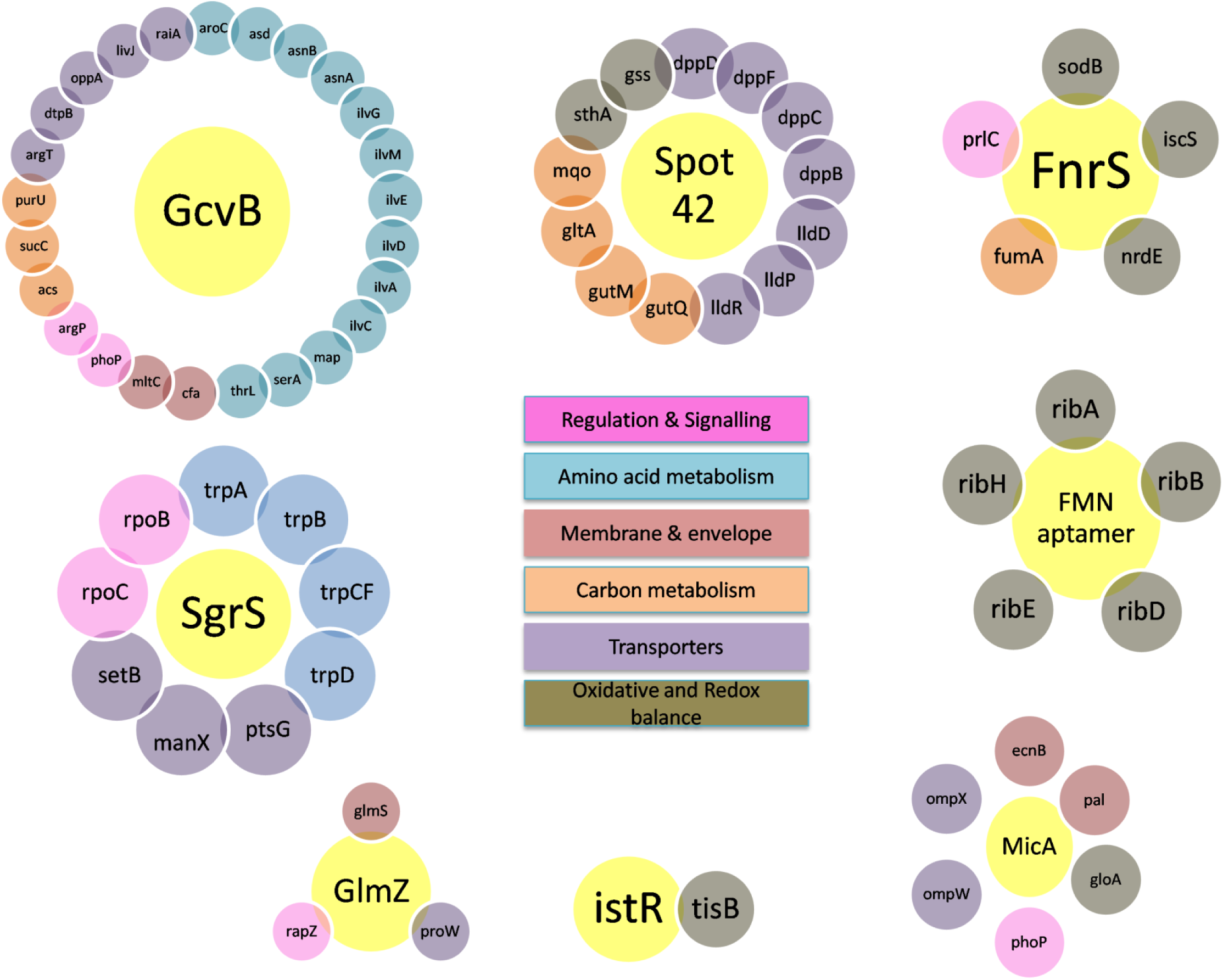
Predicted regulatory interactions between small RNAs (sRNAs) and target genes in *Klebsiella* sp. SG01 during ciprofloxacin (CIP) stress. This network diagram depicts predicted sRNA–mRNA interactions, with target genes color-coded according to associated metabolic pathways, including carbon and amino acid metabolism, transport systems, redox regulation, and general stress response. Interactions were predicted using TargetRNA3 and further supported by literature-based evidence. The figure highlights stress-induced regulatory reprogramming mediated by sRNAs, facilitating adaptive responses under CIP exposure.

The sRNA *spot42* was specifically predicted to target the Si-specific NAD(P)(+) transhydrogenase, several sugar transporter genes (e.g., *dppD, lldP*), and tricarboxylic acid cycle (TCA) enzymes, indicating roles in redox balance and carbon assimilation. By interacting with targets like *argT*, *dtpB*, and *oppA*, *gcvB* was connected to the inhibition of ABC transporters and amino acid permeases (see Supplementary File 1).

Additionally, IntaRNA analysis was used to improve the resolution of functional predictions. With the use of this tool, sRNA-mRNA base-pairing energetics could be modeled, and interaction energies between -11 and -29 kcal/mol were obtained. For instance, under low GlcN6P conditions, *glmZ* improved the translation of *glmS* by targeting its 5′ UTR (–23.8 kcal/mol). The sequence *purR* was bound by *sgrS* in the CDS (+837 nt), indicating purine biosynthesis suppression. Similarly, *fnrS* targeted *sodB*, which probably decreased superoxide dismutase activity and encouraged the buildup of reactive oxygen species (ROS), whereas *gcvB* suppressed *gdhA* and *argT* through internal CDS interactions.

Other interactions included *istR*-mediated suppression of *tisB* toxin peptide expression (+108 nt), FMN aptamer–*ribD* interaction consistent with riboswitch-induced transcriptional attenuation, and *micA* repression of *phoP* translation via 5′ UTR pairing (–29.85 kcal/mol). Table 2 provides a summary of these interactions, while Figure 3 illustrates them.

**Figure 3.**
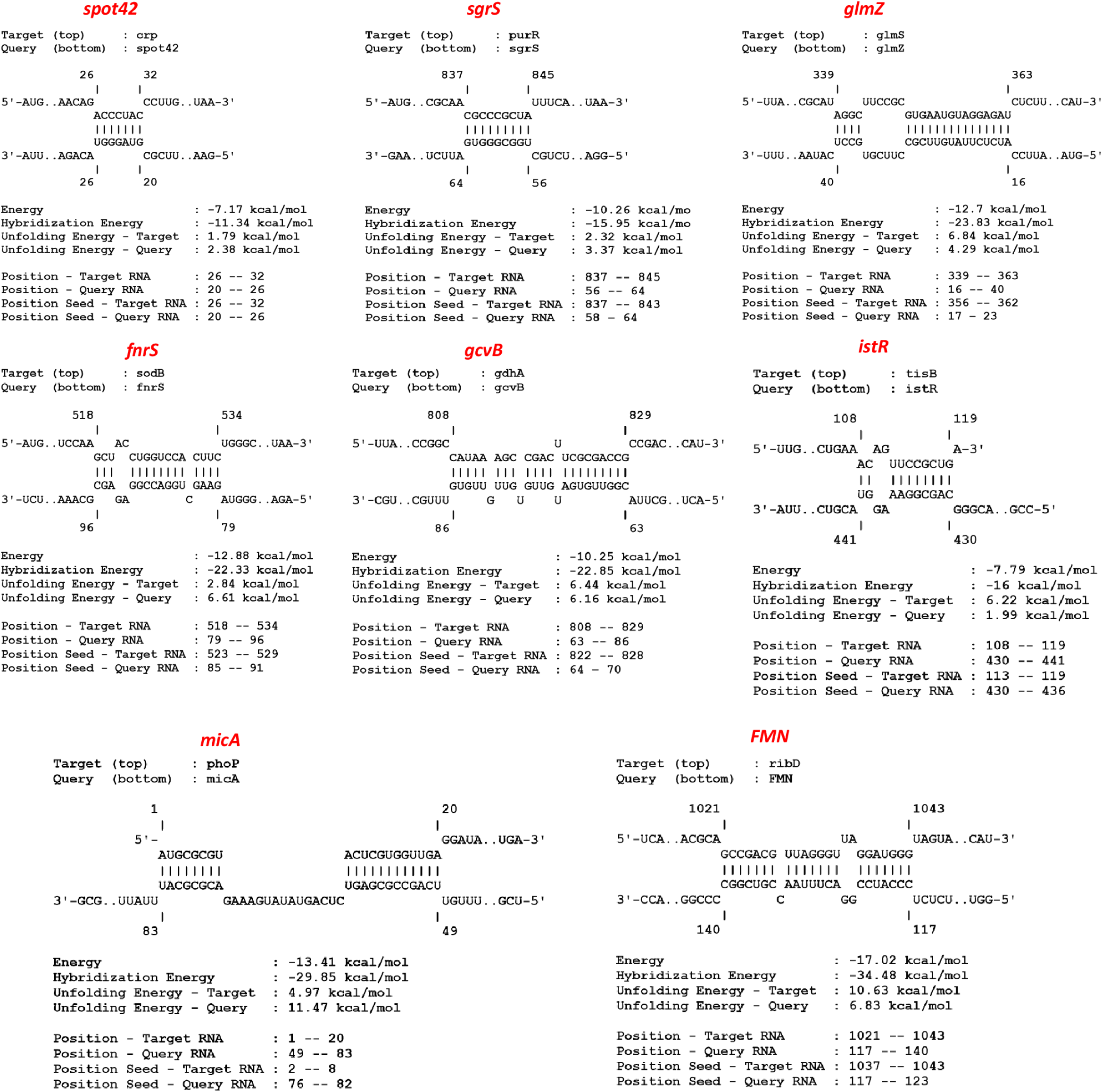
Predicted interactions of sRNAs with the regulatory regions of their mRNA targets as analyzed using IntaRNA. This figure shows the base-pairing interactions between sRNAs and the upstream regulatory regions of their respective mRNA targets, as predicted by IntaRNA. The visualization emphasizes potential post-transcriptional regulatory mechanisms involving direct binding to 5’ UTRs or proximal CDS regions.

**Table 2.**
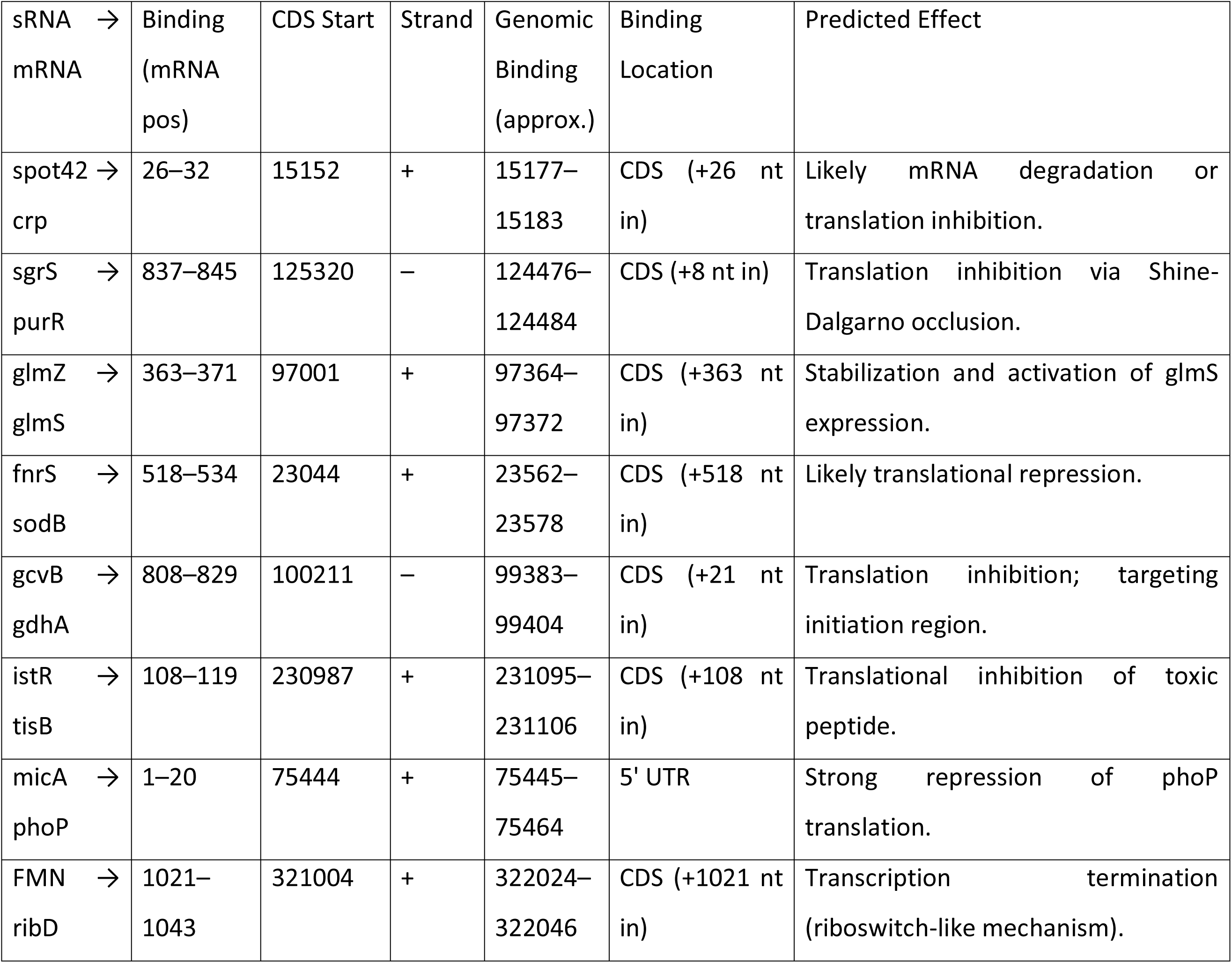
Predicted interactions between selected small RNAs (sRNAs) and their mRNA targets in *Klebsiella* sp. SG01. The table summarizes sRNA-mRNA interactions, including binding sites, coding sequence (CDS) context, strand orientation, approximate genomic binding coordinates, and anticipated regulatory outcomes. Binding locations are mapped relative to the CDS start sites, indicating whether interactions occur within CDS or untranslated regions (UTRs), and specifying the likely mode of post-transcriptional regulation.

### 2.4. Integration of Metabolomics and Transcriptomics: Evidence for Antibiotic-Derived Metabolism

Metabolomic profiling revealed the accumulation of short-chain organic acids, such as potassium propionate, succinate, and acetic acid esters in *Klebsiella* sp. SG01 cells treated with ciprofloxacin. These substances are intermediates of propionate pathway and tricarboxylic acid (TCA) cycle. Their appearance clearly points to ciprofloxacin’s oxidative breakdown and subsequent entry into key metabolic pathways.

Such catabolites most likely resemble a metabolic state that is rich in glucose. The observed transcriptomic upregulation of *spot42*, a sRNA that stabilizes in the presence of glucose or similar carbon sources, lends credence to this. This is further supported by the downregulation of CRP, a cAMP-responsive transcriptional regulator that is activated during carbon starvation.

Interestingly, hypoxanthine, a downstream purine catabolite, remained in both the log and lag phases while AMP was only found in the lag phase. This pattern implies that early purine biosynthesis was triggered in the ciprofloxacin response, which was followed by a rapid turnover or utilization of nucleotides during active growth.

A sRNA, *sgrS*, that represses *purR*, a master repressor of purine biosynthesis genes, was found to be upregulated in transcriptomic data. This likely derepresses the purine operon and facilitates the synthesis of AMP and IMP in the early adaptation to stress. Continuous detection of hypoxanthine implies an active purine salvage or catabolism for nucleotide recycling and redox buffering.

So, we would like to assert that *sgrS* attenuated antibiotic stress in addition to sugar-phosphate stress, and reprogrammed purine metabolism to modulate metabolic adaptation and survival upon antibiotic exposure.

### 2.5. Envelope and Oxidative Stress Adaptation: Roles of MicA, FnrS, and IstR

A sRNA, *micA*, regulated by σ^E^, was significantly upregulated under ciprofloxacin stress in *Klebsiella* sp. SG01. Another sRNA, *micA* is known to repress *phoP*, which encodes the response regulator of the PhoPQ two-component system. In this study, *micA* was predicted to bind the 5′ untranslated region (UTR) of *phoP* with high affinity (ΔG = –29.85 kcal/mol), potentially blocking ribosome access and promoting mRNA degradation. Since PhoPQ regulates outer membrane modifications, magnesium transport, and antimicrobial peptide resistance, its repression suggests a mechanism for membrane remodeling under antibiotic pressure.

RNA chaperone expression also showed a more general regulatory shift, with *proQ* and *csrA* both upregulated and Hfq transcripts downregulated. With ramifications for changed sRNA stability and interaction networks, this points to a potential shift in the dominant sRNA-binding proteins under stress.

Exposure to CIP significantly increased the expression of *fnrS*, a small RNA linked to adaptation to anaerobic and oxidative stress. With a ΔG of –22.33 kcal/mol, it was predicted to bind within the coding sequence of *sodB*, which codes for superoxide dismutase. Reactive oxygen species (ROS), which are essential for the oxidative breakdown of ciprofloxacin, are probably encouraged to accumulate when *sodB* is downregulated. These findings are in line with earlier research that connected *fnrS* to the increase in ROS facilitated by antibiotics.

Toxin-antitoxin systems are also activated by elevated ROS. Under stress, growth arrest and membrane depolarization are brought on by the SOS-induced peptide TisB. But the sRNA *istR*, which was upregulated in the cells treated with CIP, binds to the *tisB* coding region (+108 nt) and probably inhibits its translation. This represents a balancing act, avoiding excessive toxicity while preserving essential ROS signaling.

Together, our findings suggest that micA, fnrS, and istR act as a coordinated team. They fine-tune the bacterial response to ciprofloxacin stress by regulating redox balance, remodeling the membrane, and ultimately controlling cell survival.

### 2.6. Summary of Regulatory Integration

The strain *Klebsiella* sp. SG01 is capable of consuming CIP (2g/L) as the sole source of carbon and energy as evidenced by increase in cell number (CFU/ml). The optimum parameters for growth and degradation of CIP has been discussed (Ganguli et al. 2024).

The complex post-transcriptional regulatory network that supports bacterial adaptation to ciprofloxacin stress is demonstrated by the coordinated actions of several sRNAs that target carbon metabolism, amino acid transport, oxidative balance, and envelope integrity. A change in the use of RNA chaperones and the involvement of metabolite-responsive riboswitches further modulate this network.

Our results, which combine transcriptomic, computational, and metabolomic analyses, demonstrate a novel mechanism by which *Klebsiella* sp. SG01 dynamically rewires gene expression to withstand and possibly metabolize ciprofloxacin. Targeting sRNA circuits in antimicrobial and bioremediation strategies is made conceptually possible by these insights, which also broaden the emerging paradigm of noncanonical antibiotic assimilation.

## 3. Discussion

### 3.1. Metabolic Rewiring via sRNAs under Ciprofloxacin-Induced Stress

The available carbon source largely shapes the metabolic landscape of bacteria, and the catabolic pathways that are active are determined by regulatory networks. Ciprofloxacin was the only source of carbon and energy in our investigation, in addition to acting as a stressor. Together with transcriptomic upregulation of *spot42* and downregulation of CRP, metabolomic evidence of propionate, succinate, and acetic esters indicates that breakdown products of CIP resemble an intracellular state rich in glucose.

When there is abundant glucose in a cell, CAMP and CRP concentrations are low and *spot42* supresses the uptake of secondary carbon sources. In the absence of glucose, CAMP binds to CRP thereby suppressing *spot42*. In our study, the upregulation of *spot42* suggests that the cell is not facing any nutritional stress with respect to carbon starvation. The CIP degraded metabolites might mimic a pseudo-glucose environment, allowing metabolic reprogramming in the cells.

Another important sRNA, *sgrS*, coordinates the stress reaction to the buildup of sugar-phosphate. It targets regulatory hubs like *purR*, activates efflux transporters (*yigL*), and suppresses glucose uptake genes (*ptsG*, *manX*). This dual function of reducing sugar stress and increasing purine availability was supported in our system by upregulation of *sgrS* and metabolite profiles.

Maintenance of the cell envelope was also affected. Low levels of glucosamine-6-phosphate (GlcN6P) might be due to the downregulation of genes (*nagA*, *nagB*) involved in N-acetylglucosamine (GlcNAc) uptake. A regulatory sRNA, *glmZ*, was upregulated, stimulating the expression of *glmS* to maintain peptidoglycan biosynthesis. When recycling pathways are inhibited, this feedback loop allows the synthesis of hexosamine precursors, suggesting post-transcriptional compensation under antibiotic stress.

Similarly, amino acid metabolism was regulated by sRNAs. Glutamate is a crucial TCA intermediate and its biosynthesis was suppressed by *gcvB*. It repressed *gdhA* and *argT*, thereby balancing the utilization of carbon and nitrogen.

### 3.2. Envelope Stress Response and Post-Transcriptional Regulation via MicA

Gram-negative bacteria rely on a complex, multi-layered outer envelope. Keeping this vital barrier functioning properly is critical, especially when environmental stress hits. The σ^E^-dependent envelope stress response is essential for detecting membrane disruptions and modifying gene expression in response. When envelope integrity is compromised, σ^E^, which is normally sequestered by the anti-sigma factor RseA, becomes active. We found that *rseA* was downregulated in this study, which probably makes it easier for σ^E^ to activate when under ciprofloxacin stress.

The sRNA micA acts as a key player downstream of the stress sensor σ^E^. It works by binding onto the start of the *phoP* messenger RNA (its 5’ untranslated region), effectively blocking the ribosome and silencing *phoP* translation. Our analysis predicts this binding is extremely stable (ΔG = -29.85 kcal/mol).

PhoP is part of the PhoPQ two-component system, a master regulator controlling critical outer membrane defenses. These include resistance to antimicrobial peptides, virulence, LPS modifications, and magnesium import. By silencing *phoP, micA* indirectly dials down this entire PhoPQ defense program. This strategic shutdown helps bacteria remodel their membrane and conserve energy, allowing them to weather the storm of antibiotic attack

This interaction highlights how global stress pathways are post-transcriptionally regulated. All non-essential envelope functions are downregulated by the *micA-phoP* regulatory circuit to maintain membrane integrity upon xenobiotic stress.

### 3.3. Oxidative Stress and Toxin Regulation through *fnrS* and *istR*

Upon exposure to ciprofloxacin, cells face oxidative stress due to the buildup of reactive oxygen specis (ROS). The sRNA *fnrS*, involved in anaerobic metabolism and response to oxidative stress is upregulated in CIP-exposed cells. We found its interaction with the superoxide dismutase *sodB* where fnrS binds to the coding region of sodB with a binding energy of –22.33 kcal/mol and represses it. This is done deliberately to allow ROS buildup within the cells, using ROS to its advantage to breakdown CIP for its consumption and survival.

However, to prevent self-damage, controlled ROS elevation needs to be strictly regulated. Increased ROS in many bacteria sets off the SOS response and activates toxin-antitoxin modules, such as the TisB toxin, which stops growth and reduces ATP levels to preserve DNA integrity. It was observed that the antisense sRNA *istR* was upregulated in our system and was expected to bind within the *tisB* coding sequence. This binding probably stops translation and stops the expression of toxins unchecked.

A multi-layered survival strategy is suggested by the equilibrium between ROS accumulation and toxin regulation. While *istR* serves as a molecular brake, limiting the synthesis of TisB to prevent excessive cellular damage, *fnrS*-mediated suppression of oxidative defenses encourages the degradation of antibiotics.

Furthermore, we identified an FMN riboswitch linked to the *ribD* operon. This riboswitch probably regulates flavin biosynthesis in response to intracellular redox changes because FMN and FAD are crucial for redox enzymes like catalases and peroxidases. During oxidative stress adaptation, its presence adds an additional layer of metabolite-responsive control.

All of these results point to a complex network of regulatory RNAs, that allow *Klebsiella* sp. SG01 to use ROS as a signal for self-preservation and as a biochemical tool for the breakdown of antibiotics.

## Conclusion

In this study, we report that *Klebsiella* sp SG01 is regulated post-transcriptionally by small regulatory RNAs (sRNAs) and riboswithes to metabolize CIP as the sole source of carbon and energy. Important sRNAs regulating purine biosynthesis, redox balance, central metabolism, and envelope integrity include *spot42*, *sgrS*, *glmZ*, *gcvB*, *fnrS*, *micA*, and *istR*. The FMN aptamer further alter gene expression by sensing intracellular metabolite levels.

Under selective pressure such as antibiotic, this regulatory framework enables judicious resource allocation by shutting down non-essential pathways. Adaptive plasticity in post-transcriptional control systems is highlighted by the observed change in RNA chaperone usage (from Hfq to ProQ and CsrA).

Our research focuses on sRNA-mediated regulation in bacteria which allows it to adapt and survive under xenobiotic stresses. It is necessary to unwind these sRNA circuits in order to completely understand bacterial physiology and stress response. Additionally, it creates opportunities for the advancement of environmental bioremediation methods and antimicrobials.

## 4. Experimental Procedures

The computational workflow for sRNA prediction from the whole genome and transcriptome analysis of *Klebsiella* sp SG01 is outlined in figure 4.

**Figure 4.**
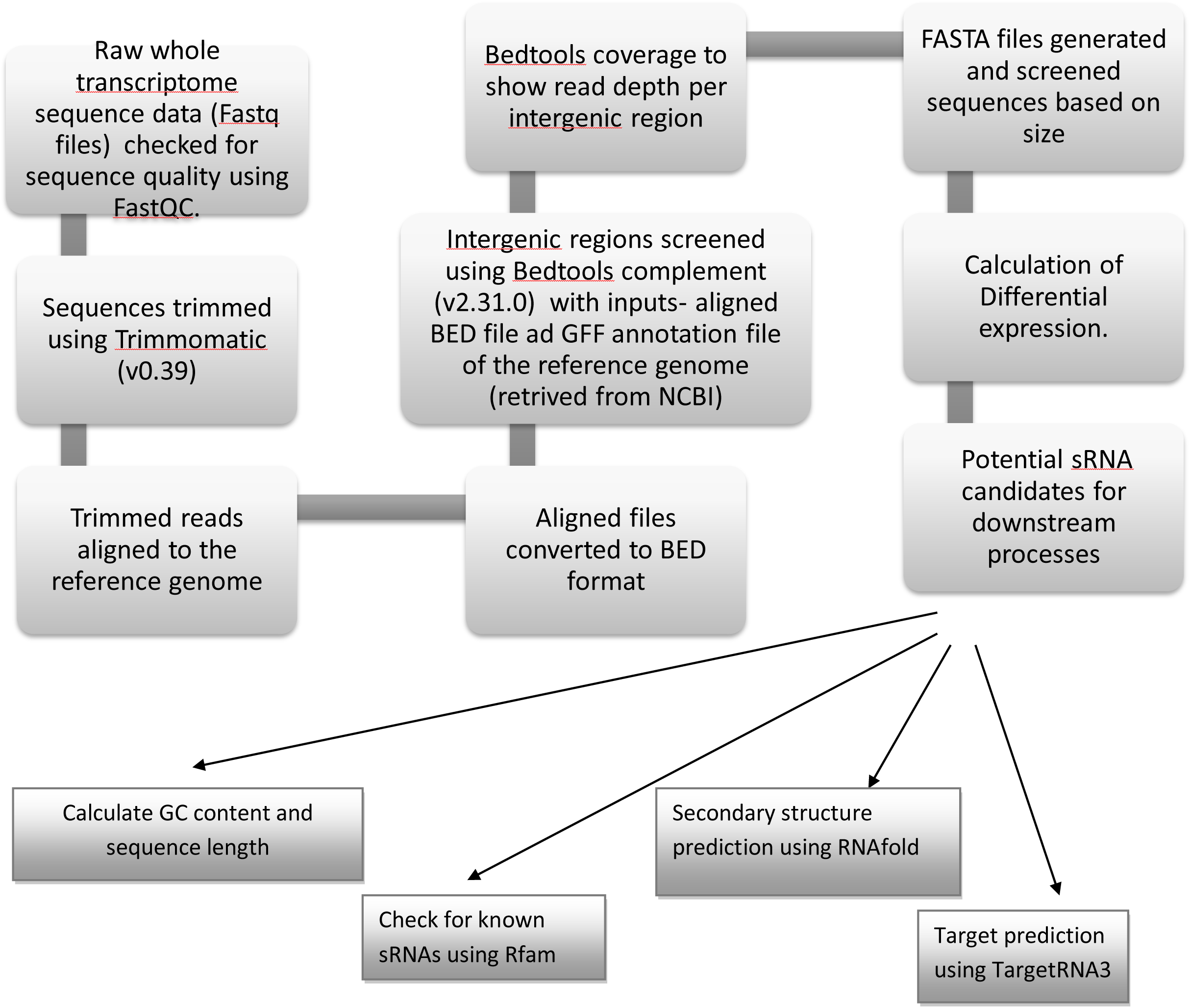
Computational workflow for the identification of small RNAs from bacterial whole-transcriptome data. This schematic diagram outlines the bioinformatics pipeline employed for sRNA identification, including transcriptome assembly, filtering, structural prediction, promoter/terminator annotation, and interaction analysis.

### 4.1. Transcriptomic Data Processing and Analysis

#### 4.1.1. Raw data processing

Paired-end FASTQ files from control and treated samples were retrieved from the National Center for Biotechnology Information (NCBI SRA SRR29374586) and Trimmomatic (v0.39) (Bolger et al. 2014) was used for quality-trimming with the following parameters: removal of Illumina adapters, sliding window trimming (4-bp window, minimum quality score 20), and retention of reads ≥15 bp. FASTQC was employed to determine the sequence quality and a Phred score>20 was used for further analysis.

#### 4.1.2. Alignment to reference genome

We aligned the trimmed reads to the Klebsiella sp. SG01 reference genome (RefSeq assembly: GCF_034387655.1) using Bowtie2 version 2.5.1 (Langmead and Salzberg, 2009). This alignment was performed in end-to-end mode using the default parameters, resulting in BAM format alignment files. The corresponding genome annotation file (in GFF3 format) was downloaded from the NCBI RefSeq database.

#### 4.1.3. Identification of Intergenic Regions

To specifically target intergenic regions, we first converted the aligned reads from the BAM files into BED format (Quinlan and Hall, 2010). We then used the bedtools complement tool (version 2.31.0, Quinlan, 2010) on these BED files, in conjunction with the genome annotation GFF3 file, to identify the non-coding regions located between annotated genes with the genome annotation file as input. Further, the BEDtools coverage by comparing aligned BAM files (Control and Treated) with the complement BED file (intergenic regions), identifying regions enriched with small RNAs. Sequences were stratified into size ranges: 15–50 nucleotide (nt), 50–250 nt, and 200–500 nt.

#### 4.1.4. Expression quantification

Reads overlapping intergenic regions were filtered by Phred score (>100). The coverage data of control and treated files were transformed to BED format. The control and treated datasets were merged, and log2-fold change (log2FC) values were calculated using

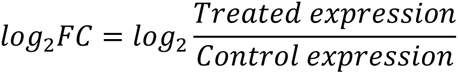

We pinpointed significantly changed regions by setting a clear threshold: only those showing a fold-change (|log2FC| > 2) were considered truly altered. To make sure our math worked reliably, especially avoiding impossible calculations like division by zero, we first added a tiny, stabilizing amount (+1) to all values before calculating the log ratios.

### 4.2. Candidate Small RNA Characterization

The small RNA candidates were compared against the Rfam database to determine homology with known non-coding RNA families. Sequences matching known ribosomal RNAs, tRNAs, or housekeeping RNAs were excluded. The minimum free energy (MFE) of candidate small RNAs was calculated using RNAfold (ViennaRNA Package v2.0) to predict their secondary structure stability.

### 4.3. Metabolite Extraction and UHPLC-MS/MS Analysis

We grew Klebsiella sp. SG01 in Minimal Salt Medium (MSM) spiked with ciprofloxacin (2 g/L). To capture different growth stages, we harvested cells during the mid-lag phase (∼6 hours) and mid-log phase (∼17 hours). Harvesting involved spinning down cells at 8000 × g for 10 minutes (4°C) followed by quickly rinsing the cell pellets with ice-cold PBS buffer and immediately halting metabolism by plunging cells into cold methanol. These quenched samples were incubated at -20°C for 1 hour. We then centrifuged them again at 14,000 × g (4°C, 20 min) to collect the metabolite-containing supernatants. These supernatants were snap-frozen in liquid nitrogen, then freeze-dried (lyophilized) under vacuum.

The dried metabolite extracts were carefully revived by dissolving them in 100 µL of LC-MS-compatible solvent with gentle vortexing. Any undissolved material was removed by a final centrifugation step (14,000 × g, 15 min), followed by filtration through a 0.22 μm membrane into LC-MS vials. Small amounts of every extract were pooled to create Quality Control (QC) samples, injected throughout the run to monitor instrument stability. Blank samples (prepared identically but without bacteria) were also analyzed to identify and subtract background signals.

An Agilent UHPLC system with a Thermo Q Exactive™ HF-X Orbitrap was used to perform UHPLC-MS/MS. A 50 × 2.1 mm Accucore HILIC column was used gradient mixing solvent A (0.1% formic acid, 10 mM ammonium acetate in 95% acetonitrile) and B (50% acetonitrile with same additives) and flow rate of 0.3 mL/min. Column was heated to 40°C and samples were kept cold at 4°C in the autosampler. The program of elution took 20 min with temperature control (column, 40°C; autosampler, 4°C). 5 µL samples were injected in random order, with QC and blank samples inserted every 5 injections. Detection of MS was done in both positive and negative ionization modes (spray voltage: 3.2 kV, capillary temp: 320°C), scanning masses from 100 to 1500 m/z. The instrument used data-dependent acquisition to automatically target the strongest signals for fragmentation (MS/MS)

### 4.4. Data Processing

Raw LC-MS data (.d files) were converted to mzML format which were processed in MZmine (Schmid et al. 2023) for peak detection, alignment, and normalization. The metabolites were identified by matching with databases like HMDB, METLIN, and LIPID MAPS (Wishart et al. 2007; Smith et al. 2005; Sud et al. 2005) based on accurate mass, retention time and spectra.

### 4.5. Regulatory Network Construction

The functional role of identified small RNAs was inferred based on their genomic location, expression patterns, and predicted target interactions. Targets of the putative sRNAs were predicted using the TargetRNA3 tool (cs.wellesley.edu/∼btjaden/TargetRNA3/) which is a web server that predicts RNA–RNA (sRNA–mRNA) interaction based on hybridization energy within a specific bacterial strain (Tjaden 2023) and also based on literature reports. Furthermore, sRNA– mRNA interaction pairs are ranked based on these hybridization energy values. A regulatory model was constructed to illustrate the influence of upregulated small RNAs on our strain’s ciprofloxacin consumption and shift in metabolism. The interaction between sRNA and mRNA was predicted using IntaRNA (http://rna.informatik.uni-freiburg.de/IntaRNA/Input.jsp;jsessionid=3F756463C9C22ED921F0734B900EE5C9) (Mann et al. 2017).

## Acknowledgments

We would like to acknowledge the University of North Bengal, Raja Rammohanpur Campus, India-734010 for their support in conducting this study. We are indebted to the Department of Biotechnology, Government of India for funding a part of our work (BT/PR40383/BCE/8/1561/2020). S.G. is thankful to the Government of West Bengal (WBP211629117511) for providing financial aid.

## Author’s Contribution

S.G. participated in designing and performing the studies, analysed data, wrote and reviewed the manuscript; R.C. conceived the idea, designed the study and wrote and reviewed the manuscript.

## Data availability

Raw sequence reads are available at SRA: <u>PRJNA931810</u> under accessions SRR24804248 for the draft genome sequence of *Klebsiella* sp. SG01 and SRR29374586 for the whole transcriptome sequence. Preprint version of the manuscript is available with doi: https://10.22541/au.174401656.60165496/v1.

## Supplementary Material

Supplemental material for this article is provided in Supplementary file 1

